# Machine learning investigation of gene expression datasets reveals *TP53* mutant-like AML with wild type *TP53* and poor prognosis

**DOI:** 10.1101/2023.02.22.529592

**Authors:** Yoonkyu Lee, Linda B. Baughn, Chad L. Myers, Zohar Sachs

**Affiliations:** Division of Hematology, Oncology and Transplantation, Department of Medicine, University of Minnesota, Minneapolis, MN USA; Bioinformatics and Computational Biology Program, University of Minnesota, Minneapolis, MN USA; Masonic Cancer Center, University of Minnesota, Minneapolis, MN USA; Division of Hematopathology, Department of Laboratory Medicine and Pathology Mayo Clinic, Rochester, Minnesota, USA; Department of Computer Science and Engineering, University of Minnesota, Minneapolis, MN USA

## Abstract

Acute myeloid leukemia (AML) with *TP53* mutations (*TP53Mut*) has poor clinical outcomes with 1-year survival rates of less than 10%. We investigated whether this AML subtype harbors a distinct gene expression profiling (GEP), what this GEP reveals about *TP53Mut* AML pathophysiology, and whether this GEP is prognostic in *TP53* wild type (*TP53WT*) AML.

We applied a supervised machine-learning approach to assess whether a unique *TP53Mut* GEP could be detected. Using the BEAT-AML dataset, we randomly divided the samples into training and testing datasets, while the TCGA dataset was reserved as a validation dataset. We trained a ridge regression machine learning model to classify *TP53Mut* and *TP53WT* cases. This model was highly accurate in distinguishing *TP53Mut* versus *TP53WT* cases in both the test and validation data sets. Additionally, we noted a cohort of *TP53WT* samples with high ridge regression scores and poor overall survival, suggesting share clinical and GEP features with *TP53Mut* AML. We defined these *TP53WT* samples as *TP53* mutant-like (*TP53Mut*-like) AMLs. We trained a second ridge regression model to specifically detect *TP53Mut*-like samples in the BEAT AML dataset and found that TCGA data also harbors *TP53Mut*-like samples. The *TP53Mut*-like samples in the TCGA also have a worse OS rate than *TP53WT* cases. Using drug sensitivity data from 122 small molecules in the BEAT AML dataset, we found *TP53Mut*-like AMLs have distinct drug sensitivity patterns compared to *TP53WT*. Finally, we identified a 25 gene signature that can identify *TP53Mut*-like cases. This signature could be used clinically to identify this novel subset of poor-prognosis AML.

## INTRODUCTION

Acute myeloid leukemia (AML) is a blood cancer with a 2-year survival rate of 30-40%^1^. *TP53* mutant (*TP53Mut*) AML is the most rapidly fatal AML subtype. Standard cytotoxic chemotherapy for AML can temporarily control the disease in most patients with *TP53Mut* AML, but *TP53Mut* AML is chemo-refractory, leading to inevitable and rapid clinical progression. The 1-year survival rate for *TP53Mut* AML is 0-10%^1–5^. A small fraction of *TP53Mut* patients stay in remission long enough to undergo allogeneic hematopoietic stem cell transplant (**HSCT**, otherwise known as bone marrow transplant), but a large proportion of these patients also inevitably relapse^6–8^. None of the FDA-approved targeted therapies for the treatment of AML have shown sustained activity in the *TP53Mut* setting. For example, the BCL2 inhibitor, venetoclax, has revolutionized the treatment of AML and is FDA approved for upfront treatment of frail and elderly AML patients^9–11^. Patients with *TP53Mut* AML have the worst treatment outcomes even when treated with venetoclax combinations with median survivals of less than 18 months^12,13^.

Gene expression profiling (GEP) has been used to interrogate the biological states of many malignancies and has been correlated with prognosis. GEP has been shown to improve prognostic classifications relative to traditional methods^14,15^ and predict therapy responses^16^. Patients whose AML expresses the leukemia stem cell (LSC) signature display a worse overall survival compared to those that do not express this signature^17,18^. These data suggest that AMLs that are more stem-like are more clinically aggressive. In other settings, GEP has been used to identify subsets of disease with unique clinical behaviors. For example, a subset of acute lymphoblastic leukemia (ALL) samples that lack the Philadelphia chromosome (Ph^-^ ALL) but share the GEP of Ph+ ALL display similar poor clinical outcomes^19,20^. In breast cancer, specimens that express a GEP resembling that of *TP53Mut* breast cancer have reduced overall survival (OS)^21^. In AML, GEP has also been used to predict chemotherapy response^16^. These findings demonstrate that GEPs can be used to infer functional states of leukemias.

Since *TP53Mut* AML is uniquely treatment refractory, we asked whether these leukemias harbor a unique GEP. We analyzed two previously published datasets that include GEP and clinical annotations of AML samples. Using a regression-based supervised machine learning model, we generated a model that can classify *TP53Mut* AML. Based on this ML model, we identified a subset of *TP53* wild type (*TP53WT*) samples in both datasets that express the *TP53Mut* GEP. These *TP53WT* cases which we call *TP53Mut*-like were found in all AML risk startification and had a worse survival rate relative to other *TP53WT* samples. Finally, we identified a 25 gene signature that can be used clinically to identify *TP53Mut*-like AML cases.

## METHODS

### Patient selection and data preprocessing

We used 2 independent data sets, BEAT AML^22^ and TCGA^23^. We acquired updated clinical data for BEAT AML^24^ and TCGA data^23^. We confirmed every sample’s ELN 2017^2^ and 2022^25^ risk stratification in the BEAT AML dataset using the updated annotations reported in ^24^. We also assigned ELN 2022 risk stratifications to each sample in the BEAT AML database and assigned ELN17 and ELN22 risk stratifications to each sample in the TCGA database. We omitted cases where insufficient information was available for FLT3-ITD allelic ratio (these cases were omitted from the ELN17 analysis, but included in the ELN22 analysis), CEBPA allelic state (these were omitted from the ELN17) and CEBPA mutation type (these were omitted from the ELN22 analysis).

All analyses were performed using R (v4.2.0) software package (R). The gene sequencing methods of publicly shared AML patients were previously described (TCGA and BEAT AML)^22,23^. We selected samples with confident *TP53* mutant calls using whole exome sequencing data from BEAT AML (n=403) and TCGA (n=178). Samples without mutational profiling were omitted from this analysis.

The raw RNA Seq read counts data were obtained from the BEAT AML^22^ and TCGA^23,26^ data. For heatmap visualization, raw read counts were normalized to counts per million (CPM) and log2 transformed using edgeR (v3.38.4) packages in R. The normalized counts were further transformed into mean-centered (Z-score) for each gene to perform clustering analyses and visualization.

For the machine learning algorithm training and testing, raw read counts from BEAT AML and TCGA data were combined and corrected for batch effects using ComBat_seq function in sva package (v3.44.0) in R with the default setting. The corrected data were log2(CPM) normalized for further analysis (**Supplemental Figure S1A**).

### Bulk gene expression profile analyses

The normalized gene expression profile data were clustered using the ComplexHeatmap (v2.12.1, Gu *Bioinformatics* 2016) package with the top 5,000 variable genes in the data. Hierarchical agglomerative clustering using the Pearson correlation coefficient as the similarity metric and the average linkage rule was used to cluster the data. Principal component analysis (PCA) was performed using the prcomp function in R.

Raw read counts of the gene expression data were used to perform differentially expression gene analysis using the edgeR package (v.3.38.4) with glmQLFit and glmQLFTest functions. GSEA was performed using the clusterProfiler package (v4.4.4) with gene sets curated from the Molecular Signatures Database (MSigDB v2022.1.Hs, https://www.gsea-msigdb.org/gsea/msigdb).

### Generation of Ridge regression classifier

To construct the *TP53Mut* AML classifier, we randomly split the BEAT AML samples into training and testing dataset by randomly dividing the dataset into 60% (n=242) and 40% (n=161) of the data. We used a logistic regression classifier with ridge regularization (ridge regression^27^) as a classifier model using the glmnet package (v4.1-6) in R^28^. A ridge regression model was generated that classified *TP53Mut* AML based on the training data using a 10-fold crossvalidation method for hyperparameter optimization. The ridge regression model was applied to the BEAT AML test and TCGA validation dataset to evaluate its performance on previously unseen expression profiles. The ridge regression model assigned *TP53Mut* Ridge Scores to each patient sample. In the BEAT AML data, *TP53WT* samples with the top 10% *TP53Mut* Ridge Scores (n=40) were defined as *TP53Mut*-like AMLs. Another ridge regression classifier was constructed to classify *TP53Mut*-like AML with the same procedure described above, but using only the *TP53Mut*-like AML as the positive examples and the *TP53WT* samples as negative examples (**Supplemental Figure S1B**). We then used the resulting *TP53Mut*-like classifier that we trained on the BEAT AML dataset to identify *TP53Mut*-like AML samples in the TCGA cohort. To assess performance of ridge regression model prediction, we used the pROC package (v1.18.0) to conduct AUROC and AUPRC curve analyses.

*TP53Mut* Ridge Scores were analyzed for some analyses. For the BEAT AML test and TCGA validation datasets, Ridge Scores were taken directly as the output of the final, optimized model obtained from the BEAT AML training set, which was fit on the entire training set using the optimal parameters obtained from 10-fold CV. For the BEAT AML training data, each patient was assigned the Ridge Score corresponding to the model derived from the single fold (out of the 10-folds used for CV) on which that patient was held out of the training.

### Identification of 25 Signature genes

To identify a subset of genes that could classify *TP53Mut*-like AML, we used elastic net regression^29^ to define a small set of genes that could achieve strong classification results in distinguishing *TP53Mut*-like cases from *TP53WT* cases. Elastic net models generally produce sparser models than ridge regression, which is well-aligned with our goal of defining a small set of genes that can support discrimination of *TP53Mut*-like cases from *TP53WT* cases. We trained the elastic net model in the BEAT AML training dataset, using 10-fold CV to select the optimal hyperparameters, and evaluated the performance of the resulting model in the BEAT AML test and TCGA validation datasets. The elastic net model resulted in a small set of nonzero coefficients (corresponding to genes) that are sufficient to classify *TP53Mut*-like samples. We iterated this process a second time where only the genes with non-zero coefficients from the first round of elastic net model fitting were included in the dataset, and we again applied elastic net to the BEAT AML training set to generate an even sparser model. This ultimately resulted in a model with 25 non-zero coefficients (the 25 signature genes). We confirmed that this small gene set was also sufficient to classify *TP53Mut*-like AML vs. *TP53 WT* AML in the BEAT AML test and TCGA validation cohorts.

### Statistical analysis

The unpaired Student’s *t*-tests were used for numerical clinical parameters (e.g., percentage of blasts in bone marrow, white blood cell counts, etc). The Benjamini-Hochberg method was used to correct for the multiple hypothesis testing and calculate false discovery rate (FDR). Fisher’s exact test was used for analysis of enrichment of cytogenetics and mutational status in BEAT AML and TCGA data. Survival analyses were conducted using Kaplan-Meier method and Cox proportional-hazards model with log-rank tests implemented in the survival (v.3.4-0) packages in R.

### BEAT AML *ex vivo* drug sensitivity data analysis

Previously published data of the *ex vivo* drug sensitivity screen of 122 small molecule inhibitors in the BEAT AML dataset^22^ was used. The unpaired Student’s *t*-test was used to calculate the statistical significance of area under the curve (AUC) differences between *TP53Mut* vs. TP53*WT* or *TP53Mut*-like vs. *TP53WT* groups. The Benjamini-Hochberg method was used to correct for the multiple hypothesis testing and calculate FDR. We z-score normalized AUC values multiplied by −1, and created a heatmap using the ComplexHeatmap package.

## RESULTS

### *TP53Mut* AML harbors a unique gene expression profile

As previously described, *TP53Mut* AML patients have poor prognosis (BEAT AML^22^: **Figure 1A** and TCGA^23,26^: **Supplemental Figure S1C**). To define the GEP of AML with *TP53Mut*, we compared the GEP of samples with *TP53Mut* to those with wild type *TP53 (TP53WT*). After correcting for batch-effects between BEAT AML and TCGA GEP data (BEAT: *TP53Mut* n=36, *TP53WT* n=367; TCGA: *TP53Mut* n=15, *TP53WT* n=163), we performed both principal component analysis (PCA) and unsupervised hierarchical clustering methods and detected no significant clustering according to *TP53* mutation status in both BEAT and TCGA cohorts (**Figure 1B-C, Supplemental Figure S1D**). We reasoned that a supervised machine learning approach may be able to learn a *TP53Mut* GEP signature, even if such a signature was not apparent from unsupervised clustering analysis. To this end, we separated the BEAT AML dataset into training and test datasets. We used a logistic regression with ridge regularization (ridge regression) as a classifier model to learn the unique GEP features that are important to define *TP53Mut* AML. We optimized and trained our ridge regression model on a training dataset (60% of the BEAT AML data) to classify *TP53Mut* and WT cases (see Methods). We evaluated our trained classifier model on a held-out test dataset (40% of the BEAT AML data) and found that this model was highly accurate (**Figure 1D**, area under the receiver operating characteristic curve (AUROC): 0.976 (93% sensitivity and 97% specificity) and area under the precision-recall curve (AUPRC): 0.887 (86% precision and 86% sensitivity)). When we applied this model to the TCGA dataset for validation, we found a similar accuracy in classifying *TP53Mut* cases (AUROC: 0.995 (100% sensitivity and 99% specificity) and AUPRC: 0.922 (94% precision and 100% sensitivity)). The ridge regression model score (*TP53Mut* Ridge Score) reflects how closely a GEP resembles that of *TP53Mut* AML GEPs. As expected, *TP53Mut* AMLs have high *TP53Mut* Ridge Scores and poor OS in both datasets (**Figure 1E**).

**Figure 1.**
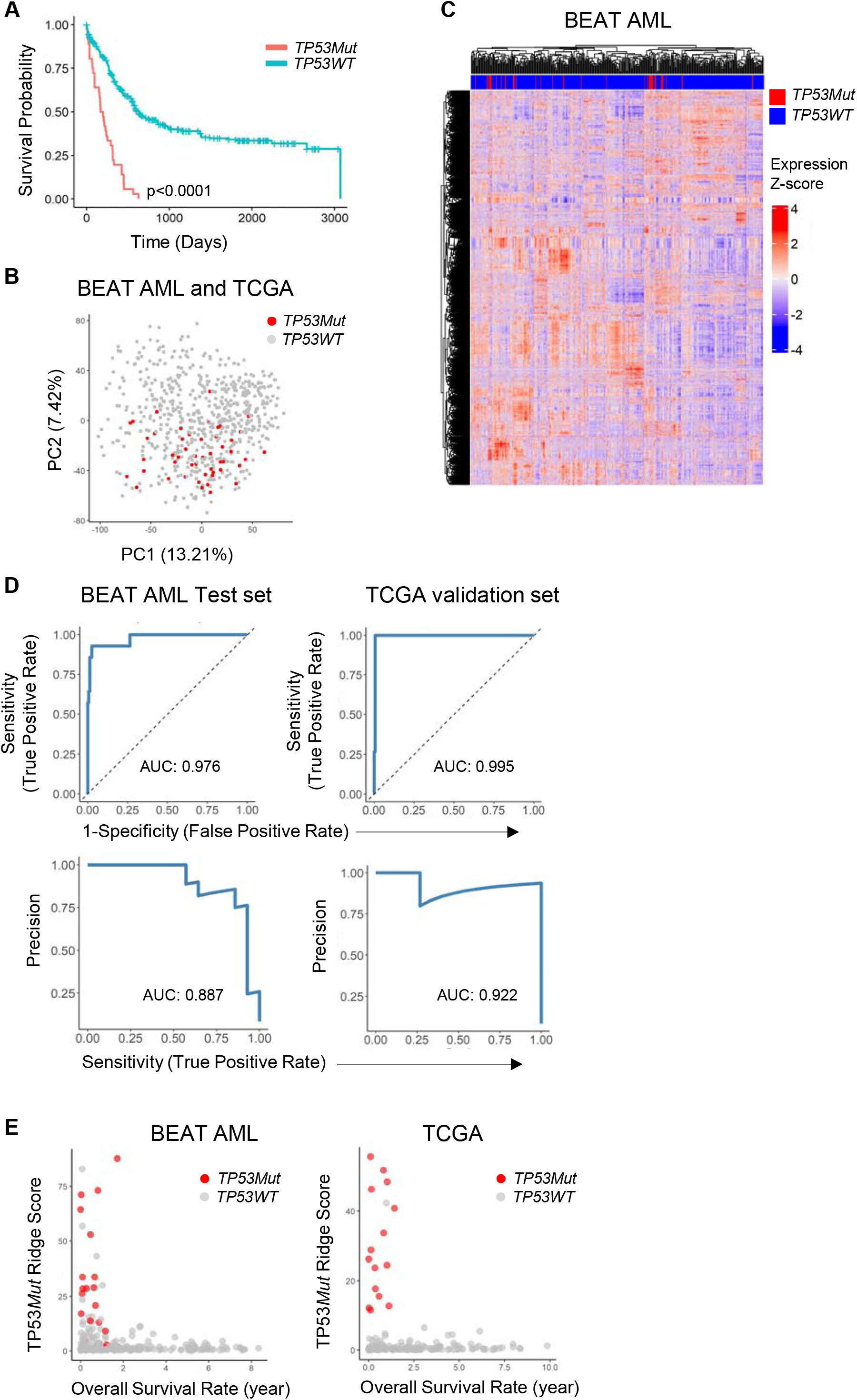
Ridge regression model classifies *TP53Mut* AML samples in BEAT AML and TCGA. (A) Kaplan-Meier estimates of overall survival (OS) curves comparing *TP53Mut* (n=36) *vs.TP53WT* (n=367) AML patients in BEAT AML data. Log-rank tests were used to calculate *P* values. (B) Principal Component Analysis (PCA) of samples in the BEAT AML and TCGA dataset after batch correction. (C) Unsupervised two-dimensional hierarchical clustering of BEAT AML samples (n=403); count per million (CPM) expression values were log2 transformed and mean-centered to generate Z-scores. (D) The performance of the ridge regression model is represented by sensitivity, specificity, and precision which are calculated by the Area Under the Receiver Operating Characteristic (AUROC) and the Area Under the Precision Recall Curve (AUPRC). (E) The *TP53Mut* Ridge Scores are plotted versus overall survival for diagnostic samples in the BEAT AML and TCGA datasets.

### A subset of *TP53WT* AML samples shares GEP features and poor clinical outcomes of *TP53Mut* AML

Strikingly, we noticed a subset of *TP53WT* cases with high *TP53Mut* Ridge Scores (representing high similarity to the *TP53Mut* GEP, **Figure 1E**). Among the samples obtained at diagnosis, these high scoring *TP53WT* cases also had low overall survival (OS) relative to *TP53WT* cases with lower scores (**Figure 2A**). In the BEAT AML dataset, we found that *TP53WT* samples in the top 10% of *TP53Mut* Ridge Scores (n=40 samples). We used only diagnostic AML samples (n=269) and diagnostic AML harbors *TP53Mut*-like AML (n=26). In the diagnostic BEAT AML, *TP53Mut*-like cases had the worse OS rates relative to the other diagnostic *TP53WT* cases (**Figure 2B**, median survival: 204 days versus 806 days, p=0.0062). We hypothesized that these diagnostic *TP53WT* cases with GEP and OS similar to *TP53Mut* AML may comprise a novel subset of AML, hereafter referred to as “*TP53Mut*-like” AML. Further, we reasoned that a machine learning could be used to specifically detect these *TP53Mut*-like cases. To test this, we trained a new complementary ridge regression model to classify these *TP53Mut*-like cases from *TP53WT* cases in the BEAT AML training data (excluding *TP53Mut cases*). We found that this new model was highly sensitive and specific in classifying held-out the test patients (AUROC:0.959 (88% sensitivity and 96% specificity) and AUPRC: 0.707 (70% precision and 88% sensitivity), **Figure 2C**). We applied this *TP53Mut*-like model to the TCGA data (which includes only diagnostic samples) and validated that high *TP53Mut*-like Ridge Scores identify a subset of *TP53WT* patients with poor OS in TCGA as well (**Figure 2D-E** and **Supplemental Figure S2A**; median survival 335 days versus 800 days, p=0.031). We visualized the TP53Mut-like cases in a PCA of BEAT AML and TCGA data and found that these cases loosely cluster together, but do not form a distinct cluster relative to the other *TP53WT* cases (**Supplemental Figure S2B**). The *TP53Mut*-like classifier was trained only on *TP53WT* cases in the BEAT AML training dataset. Interestingly, the *TP53Mut*-like ridge model classifier assigned high scores to the *TP53Mut* cases as well (**Figure 2A, D**) suggesting that this classifier specifically detects similarity to *TP53Mut* AML. The fraction of *TP53Mut*-like cases in each dataset (10-13%) was larger than the fraction of *TP53Mut* cases in each dataset (7-9%, **Figure 2F**) demonstrating that this novel subset accounts for a significant fraction of AML cases. Deletion of the *TP53* locus, in the absence of *TP53Mut* is quite rare (approximately 2%, **Supplemental Table S1**) in AML^30,31^ and equally rare in MDS (0.3-2.4%, **Supplemental Table S1**)^30,32,33^. Biallelic deletions in *TP53* have not been detected in large AML studies^30^. These observations effectively exclude a biallelic *TP53* deletion as a major contributor to the *TP53Mut*-like phenotype. We asked whether monoallelic deletions, in the absence of a *TP53Mut*, could account for the *TP53Mut*-like phenotype. The BEAT AML dataset does not include copy number assessment so we evaluated the frequency of 17p alterations (17pAlt) as a surrogate. We found that 15% of the *TP53Mut*-like cases harbor 17pAlts (6 of 40 cases, **Figure 2G**). We reviewed the cytogenetic and copy number data in the TCGA dataset and found that 22% of the *T53Mut*-like cases harbor deletions in the *TP53* locus (per copy number array and/or 17pAlt, 5 of 23 cases, **Figure 2H**). Notably, three *TP53WT* cases also harbor *TP53* locus deletions in the TCGA dataset. These data suggest that loss of the *TP53* locus is more common in *TP53Mut*-like cases but does not account for the majority of those cases and is not sufficient to induce a *TP53Mut*-like phenotype. These results define *TP53Mut*-like AML as a novel subgroup of *TP53WT* AML with poor clinical outcomes.

**Figure 2.**
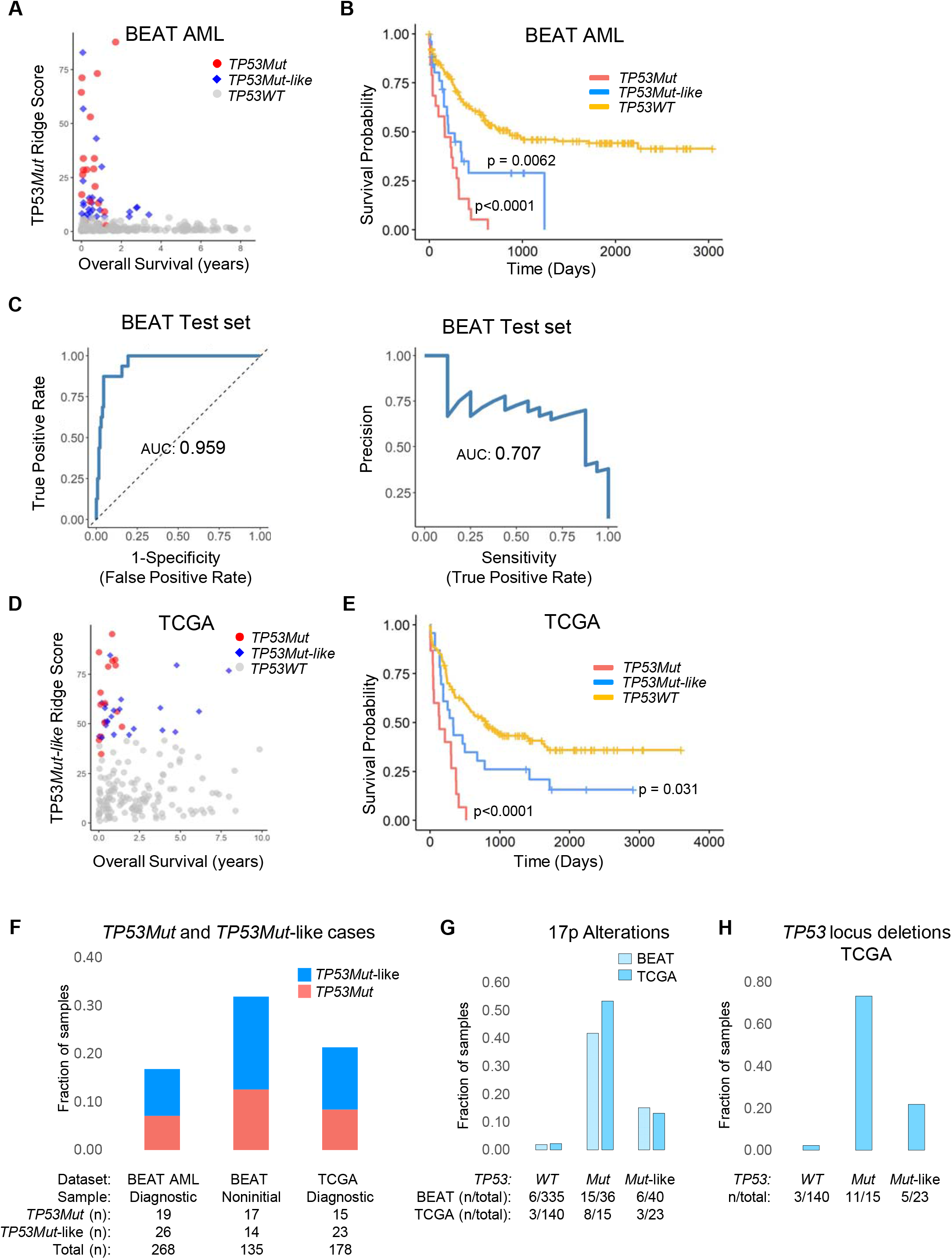
*TP53Mut*-like AML: a subset of *TP53WT* AMLs that share GEP features and poor clinical outcomes with *TP53Mut* AML. (A) *TP53Mut* Ridge Scores are plotted versus overall survival in the diagnostic samples in BEAT AML dataset. (B) Kaplan-Meier survival curves of diagnostic samples in the BEAT AML dataset. (C) *TP53Mut*-like ridge regression model performance in the BEAT AML test dataset is represented by sensitivity, specificity, and precision which are calculated by the Area Under the Receiver Operating Characteristic (AUROC) and the Area Under the Precision Recall Curve (AUPRC). (D) *TP53Mut*-like Ridge Scores are plotted versus survival in the TCGA validation dataset. (E) Kaplan-Meier survival curves of samples in the TCGA AML dataset. (B-E): *P* values reflect comparison of *TP53Mut* or *TP53Mut*-like to *TP53WT* samples. Median survival: BEAT AML *TP53Mut*-like: 204 days, BEAT AML *TP53WT:* 806 days, p = 0.0062; TCGA *TP53Mut*-like: 335 days, TCGA *TP53WT:* 800 days, p = 0.031. BEAT AML: *TP53Mut* n=36 (19 diagnostic samples), *TP53Mut*-like n=40 (26 diagnostic samples) and *TP53WT* n=335 (223 diagnostic samples. TCGA: *TP53Mut* n=15, *TP53Mut*-like n=23 and *TP53WT* n = 140. (F) The fraction of *TP53Mut* and *TP53Mut*-like AMLs in BEAT AML and TCGA data. (G-H): Fractions of all samples in each category that harbor (G) 17pAlt or **H.** *TP53* locus deletions.

### *TP53Mut*-like AML share poor clinical parameters with *TP53Mut* AML

We compared the clinical and molecular features of *TP53Mut, TP53Mut*-like and *TP53WT* cases. In comparison to the *TP53WT* samples, *TP53Mut* and *TP53Mut*-like patients have lower bone marrow (BM) blasts, white blood cell counts (WBCs), and peripheral blood (PB) blast percentages in the BEAT AML dataset (**Figure 3A, B and Supplemental Table S2**). Notably, the samples in the TCGA dataset display the same trend, but lack statistical significance, likely due to a smaller sample size (**Supplemental Figure S3A, B and Supplemental Table S2**). Like *TP53Mut* AML patients, patients with *TP53Mut*-like AML are older at diagnosis than other *TP53WT* patients in both datasets (**Figure 3C**, **Supplemental Figure S3C and Supplemental Table S2**). In addition, *TP53Mut* and *TP53Mut*-like samples have higher LSC17 scores^17^ relative to *TP53WT* cases (p<0.0001) in both datasets, indicating higher expression of leukemia stem cell-associated genes (**Figure 3D**, **Supplemental Figure S3D and Supplemental Table S2**). These differences in laboratory parameters between *TP53Mut* and *TP53WT* AMLs were consistent with prior reports^34,35^. Together, these data suggest that the *TP53Mut*-like AML resembles the clinical and biological characteristics of *TP53Mut* AML and is distinct from *TP53WT* AML.

**Figure 3.**
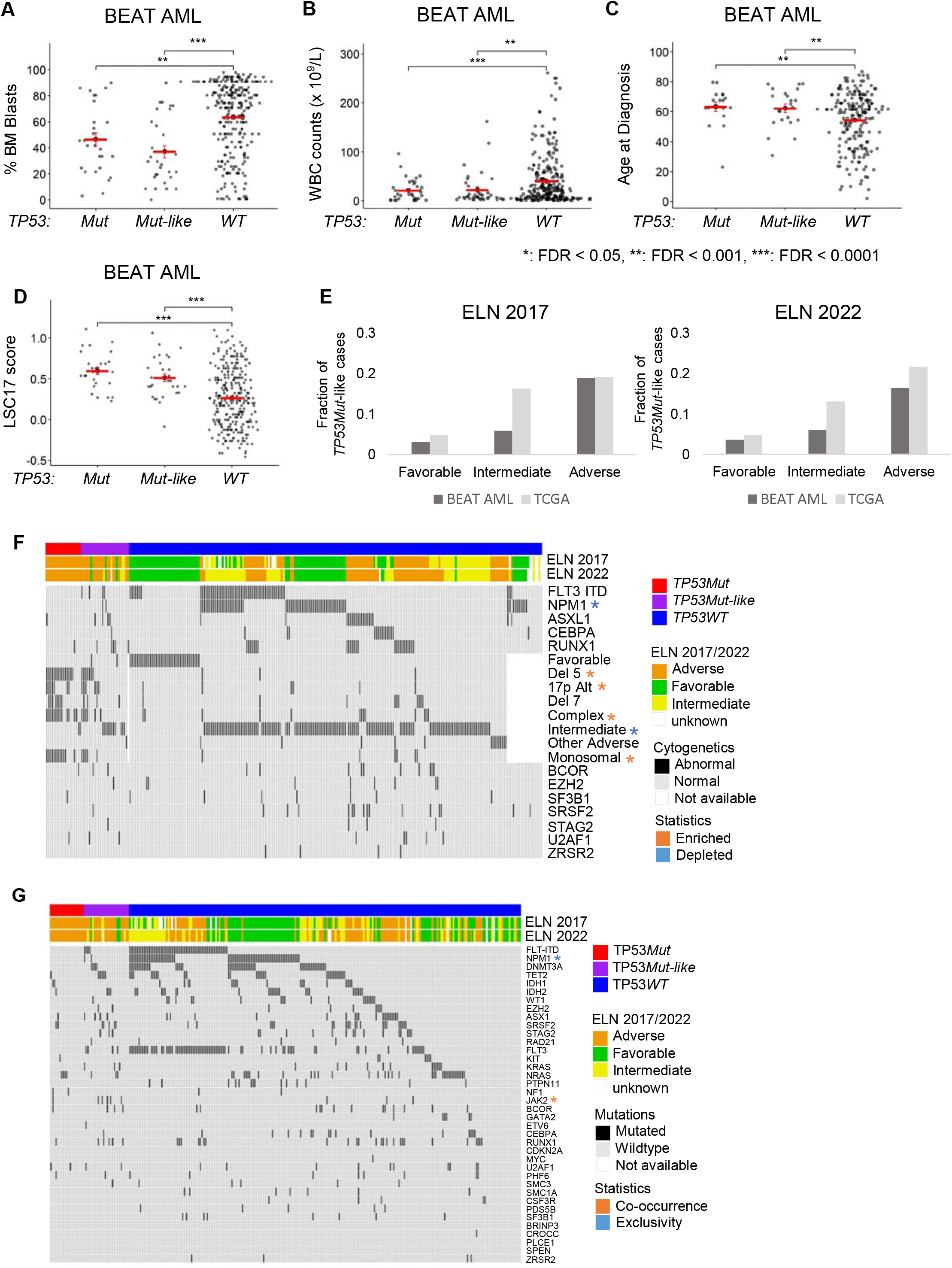
*TP53Mut*-like AML share clinical parameters with *TP53Mut* AML. Percentages of (A) bone marrow blasts, (B) white blood cell counts, (C) age, and (D) LSC17 scores were plotted for each TP53*Mut*, TP53*Mut*-like, and TP53*WT* AML diagnostic samples in the BEAT AML dataset. Horizontal red bars indicate the mean values. Unpaired Student *t*-test was used to calculate *P* values for each comparison. Benjamini-Hochberg method was used to correct *P* values for multiple hypothesis testing and to calculate the false discovery rate (FDR) (E) Fraction of diagnostic cases that are *TP53Mut*-like in each ELN risk category. (F) Cytogenetic landscape of TP53*Mut*, *TP53Mut*-like and TP53*WT* in BEAT AML diagnostic samples. (G) Mutational landscape of most frequently mutated genes of TP53*Mut*, TP53*Mut*-like and TP53*WT* in BEAT AML diagnostic samples. (F-G): Asterisks represent abnormalities whose enrichment/co-occurrence (orange asterisks) or depletion/exclusivity (blue asterisks) is statistically significant (P<0.05).

Next, we compared European Leukemia Net (ELN) risk stratifications of diagnostic *TP53Mut*-like and *TP53WT* cases. *TP53Mut* is classified as adverse risk by the ELN classifications of 2017 and 2022^2,25^. We found *TP53Mut*-like cases in all three ELN categories in both the BEAT AML and TCGA cohorts. *TP53Mut*-like cases represented 3.1-4.8% of the favorable risk cases in both datasets (BEAT AML: 3 cases; 3.1% per ELN 2017 and 3.6% per ELN 2022; TCGA: 3 cases: 4.8% per ELN 2017 and 4.8% per ELN 2022, **Figure 3E, Supplemental Figure S3E and Table S3**). As expected, *TP53Mut*-like AMLs were largely enriched in the adverse risk category (BEAT AML 20 cases; 19% per ELN 2017 and 18 cases; 16% per ELN 2022; TCGA: 12 cases; 19% per ELN 2017 and 13 cases; 22% per ELN 2022). Next, we analyzed the hazard ratio for survival in each ELN risk group found that *TP53Mut*-like cases have a trend towards inferior survival in both the favorable and adverse risk categories but the number of cases was small and this trend did not meet statistical significance (**Supplemental Figure S3E and Table S3**).

Next, we investigated whether the *TP53Mut*-like AML samples harbor unique cytogenetic or mutational features compare to *TP53WT* AMLs. We found that *TP53Mut*-like AMLs resemble *TP53Mut* AML in that they were enriched for deletion 5, 17pAlt, Complex cytogenetic abnormalities and monosomy and were depleted for Intermediate cytogenetic abnormalities in BEAT AML data (**Figure 3F and Supplemental Table S4**). In TCGA, *TP53Mut*-like AML showed enrichment of 17pAlt and deletion 7, and depletion of Favorable cytogenetics (**Supplemental Figure S3F and Supplemental Table S4**). We found some enrichment of abnormalities in mutational profiling, but these enriched abnormalities were specific to each database. *TP53Mut*-like AML harbors unique mutational profiles with cooccurrence of JAK2, and exclusivity of NPM1 in BEAT AML dataset. In TCGA, *TP53Mut*-like AML has co-occurrence of KRAS and RUNX1 mutations, and exclusivity of FLT3 mutation (**Figure 3G** and **Supplemental Figure S3G** and **Supplemental Table S5**).

### *TP53Mut*-like AML shares *ex vivo* drug resistance profiles with *TP53Mut* AML

*TP53Mut* AML is uniquely treatment refractory and patients with this AML subtype display the worst outcomes in clinical trials^12,13^. We asked whether *TP53Mut*-like AML samples might also be drug resistant. We analyzed the *ex vivo* drug-response data reported in the BEAT-AML dataset which included 122 small molecule inhibitors. When compared to the *TP53WT* cases, the *TP53Mut*-like cases resembled *TP53Mut* cases and showed resistance to a majority of the drugs. (**Figure 4A and Supplemental Figure S4A**). Next, we compared the drug sensitivity values between the *TP53Mut*-like and *TP53WT* cases. There was a trend towards increased sensitivity to 2 drugs (flavopiridol and panobinostat) but this trend did not achieve statistical significance (**Figure 4B**). Notably, flavopiridiol was also recognized as having *ex-vivo* activity in chemorefractory samples^16^. Like *TP53Mut* AML, the *TP53Mut*-like samples showed the highest resistance to venetoclax (**Figure 4C**), suggesting that these patients might benefit from alternate therapies. This result demonstrated that *TP53Mut*-like AML have drug sensitivity patterns that resemble those of *TP53Mut* AML and are different from *TP53WT* AMLs.

**Figure 4.**
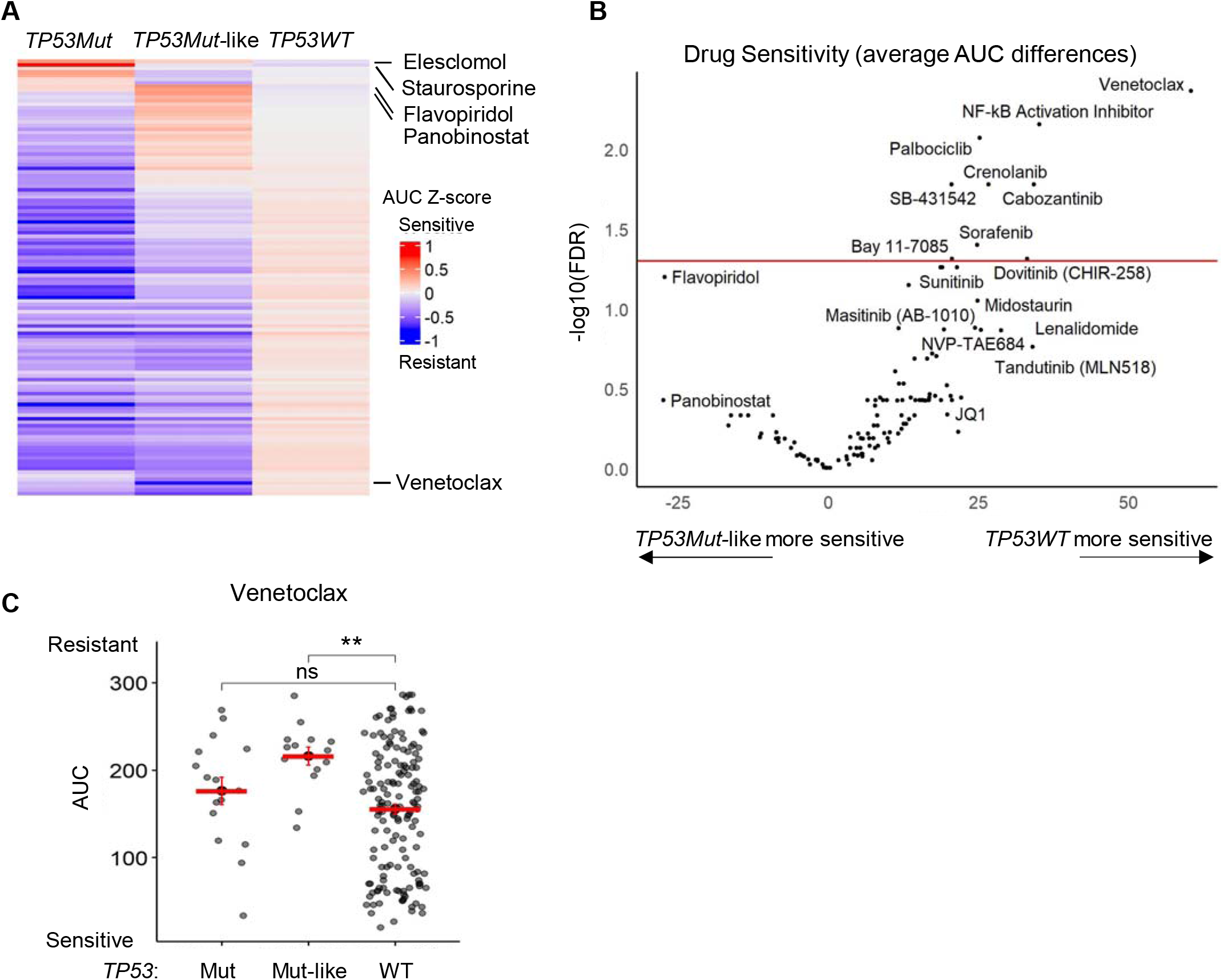
*TP53Mut*-like AML samples are drug-resistant in *ex vivo* functional drug sensitivity screening. (A) *Ex vivo* drug sensitivity data generated from 122 small molecule inhibitors in BEAT AML data. The area-under-the curve (AUC) values were Z-score transformed and multiplied by −1 (AUC Z-score). (B) Unpaired Student *t*-test was used to compare the average differences in AUC drug responses between TP53*Mut*-like and TP53*WT* samples. Multiple hypothesis testing was corrected and FDR was calculated using the Benjamini-Hochberg method. Red line indicates cutoff of FDR=0.05. (C) Venetoclax sensitivity data comparing *TP53Mut, TP53Mut*-like and *TP53WT*.

Interestingly, the resistance profile of *TP53Mut*-like samples did not fully recapitulate that of the *TP53Mut* samples. Additionally, the *TP53Mut*-like samples displayed increased resistance to venetoclax, in comparison to *TP53Mut* samples. However, none of the differences between *TP53Mut* and *TP53Mut*-like drug responses achieved statistical significance (**Supplemental Figure S4B**).

### *TP53Mut* AMLs harbor unique pathways

To identify biological functions that are specific to *TP53Mut* AML, we performed differentially expressed gene (DEG) analysis to compare *TP53Mut* and *TP53WT* AMLs. We analyzed the BEAT AML and TCGA separately and found 2460 overlapping genes (1339 upregulated and 1121 downregulated genes) that are concordantly and differentially expressed with FDR<0.05 in both datasets (**Supplemental Table S6**). Next, we repeated this analysis with the same criteria for *TP53Mut*-like AML, relative to *TP53WT* cases and found 3009 overlapping genes (1229 upregulated and 1780 downregulated genes, **Supplemental Table S6**). *TP53Mut* and *TP53Mut*-like AML share 827 genes that were concordantly and differentially expressed when each group is compared to *TP53WT* AMLs (FDR<0.05 in both BEAT AML and TCGA in each comparison) (**Figure 5A and Supplemental Figure S5**). Previous work demonstrated that some *TP53WT* AML samples express high levels of MDM2 protein, the negative regulator of p53^36^. These MDM2^High^ cases display the same poor overall survival outcomes as *TP53Mut* cases. Even though MDM2 is canonically regulated at the protein level, we found a trend towards elevated *MDM2* transcripts in *TP53Mut*-like cases, relative to *TP53Mut* and *TP53WT* cases suggesting that the *TP53Mut*-like phenotype might correlate with MDM2 activity (**Supplemental Figure S5B-C**. Interestingly, *CDKN1A* (whose transcription is induced by wildtype p53) is elevated in some *TP53Mut*-like cases suggesting that some p53 functions might be retained in these cases.

**Figure 5.**
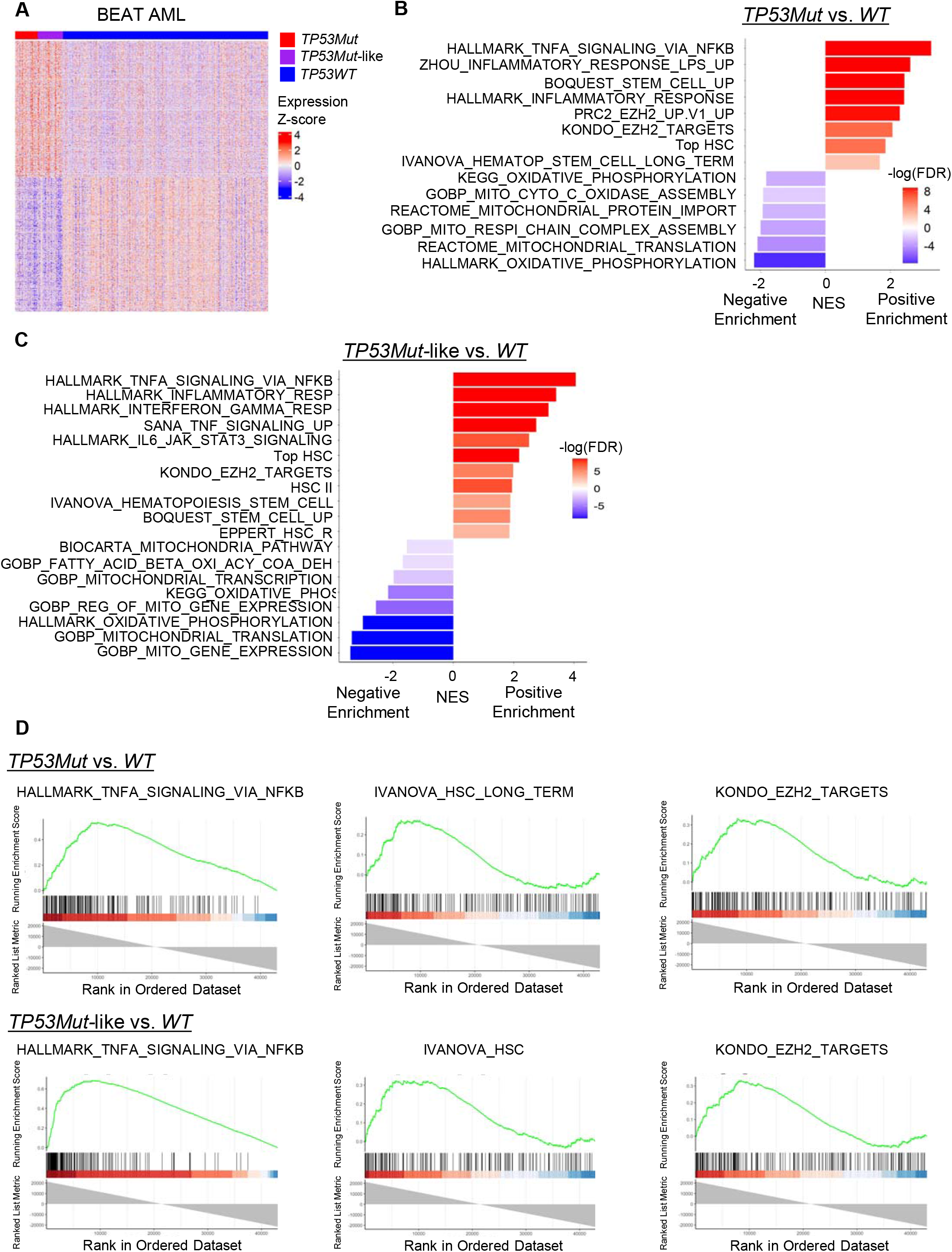
*TP53Mut*-like and *TP53Mut* AML GEP reveals unique biological pathways. (A) Heatmap of 827 differentially expressed genes (DEG) that are shared between *TP53Mut*-like and *TP53Mut* samples, in comparison to WT AMLs in both the BEAT AML and TCGA datasets. CPM expression values were log2 transformed and mean-centered to generate Z-scores. BEAT AML data is shown. (B-C): GSEA was performed to compare (B) *TP53Mut* and *TP53WT* samples, and (C) *TP53Mut*-like and *TP53WT* samples. Gene sets displayed are those that are significantly enriched in both datasets, based on concordant normalized enrichment scores (NES) and FDR values<0.05 in both datasets independently. (D) Representative GSEA plots.

We performed gene set enrichment analysis (GSEA) comparing *TP53Mut* to *TP53WT* and *TP53Mut*-like to *TP53WT* (each comparison was performed separately in each dataset). For each comparison, we identified overlapping gene sets as the gene sets with significant enrichment (FDR<0.05) and concordantly enriched (NES in the same direction) in both datasets (representative data in **Figure 5B-D**, data in **Supplemental Tables S7**). Notably, both *TP53Mut* and *TP53Mut*-like AMLs were strongly enriched with NFκB pathways, Inflammatory pathways, EZH2 targets genes and stem cell related gene sets. In contrast, *TP53Mut* and *TP53Mut*-like AMLs displayed negative enrichment (downregulation) of oxidative phosphorylation and mitochondrial pathways.

### 25 signature genes define *TP53Mut*-like AML cases

We interrogated the GEP data to identify markers of *TP53Mut* and *TP53Mut*-like cases. In comparing *TP53Mut* and *TP53WT* cases, we identified 13 genes that encode cell surface markers that were significantly and concordantly differentially expressed in both datasets (**Figure 6A**, **Supplemental Figure S6A, Supplemental Table S6**, FDR<0.05 in each dataset). In comparing *TP53Mut*-like to *TP53WT* cases, we identified 16 cell surface marker-encoding genes that were significantly and concordantly differentially expressed in both datasets (**Figure 6B, Supplemental Figure S6B, Supplemental Table S6**, FDR<0.05 in each dataset). Among these genes, five genes were concordantly differentially expressed in both comparisons. These cell surface markers offer potential therapy targets for these AML subsets.

**Figure 6.**
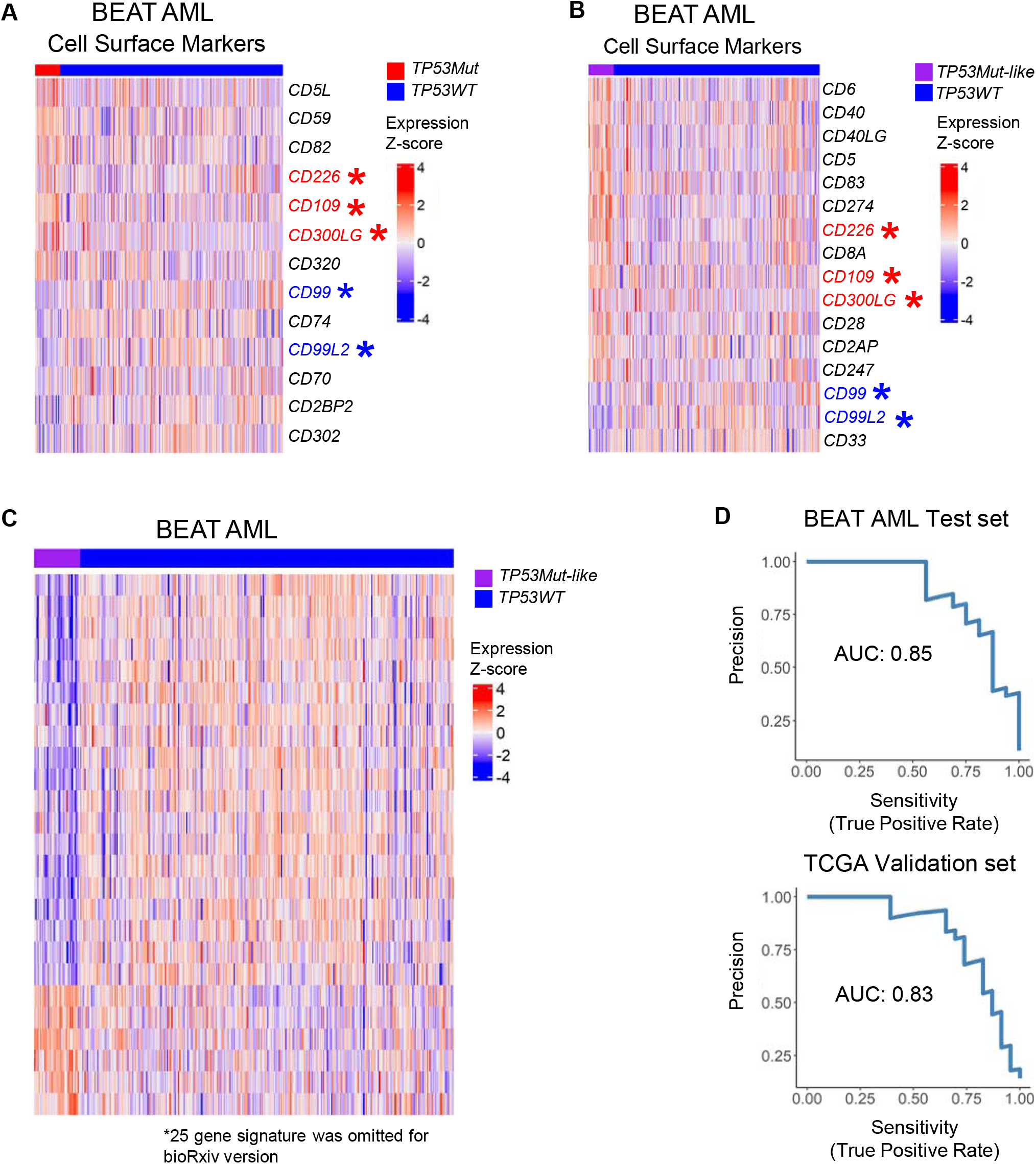
25 gene signature defines *TP53Mut*-like AML. (A-B): Genes encoding cell surface markers that are differentially expressed between (A) *TP53Mut* and *TP53WT*, and (B) *TP53Mut*-like and *TP53WT* samples. BEAT AML data. Genes encoding cell surface markers that are displayed are those concordantly differentially expressed in both the BEAT AML and TCGA datasets with an FDR<0.05 in each dataset. Data is displayed as log2 transformed count per million (CPM) expression values that were mean-centered. Genes that are concordantly shared between *TP53Mut* and *TP53Mut*-like (A and B) are the marked red (up-regulated) and blue (down-regulated) asterisk. BEAT AML data is shown. (C) Expression values of *TP53Mut*-like signature genes in BEAT AML samples. CPM values were log2 transformed and Z-score converted. (D) 25 gene signature performance in the BEAT AML test and TCGA validation datasets are evaluated using ridge regression and represented by sensitivity and precision which are calculated by the AUPRC.

Next, we asked whether a concise gene signature could be used to identify *TP53Mut*-like AML. Such a signature could be developed into a clinical assay to identify these patients. To identify a subset of genes that can classify TP53*Mut*-like AML cases, we used elastic net regression, which results in sparser models than our original ridge regression approach^29^. Using multiple rounds of elastic net model optimization (see Methods), we identified a sparse model involving 25 genes that supported high classification accuracy for *TP53Mut*-like AML vs. *TP53WT* in both datasets using elastic net or ridge regression (**Figure 6C-D, Supplemental Figure S6C-D)**.

## DISCUSSION

Using publicly available GEP datasets and a regression-based machine learning classifier, we defined a unique *TP53Mut* AML GEP and *TP53Mut*-like AML, a novel subtype of *TP53WT* AML that phenocopies *TP53Mut* AML. Notably, this subset is imperceptible using traditional clustering methods and demonstrates the power of machine learning approaches to discover new biology. *TP53Mut*-like AMLs were found in both BEAT AML and TCGA and had inferior OS rates compared to *TP53WT* AMLs. In comparison to *TP53WT* AMLs, *TP53Mut*-like AMLs had distinct clinical parameters, resembling those of *TP53Mut* AML. *TP53Mut*-like AMLs also share important biological pathways with *TP53Mut* AML. Using the BEAT AML *ex vivo* drug sensitivity data, we also demonstrated *TP53Mut*-like AMLs had unique drug sensitivity patterns and significant resistance to venetoclax. Finally, we discovered a 25 gene signature that can be used to identify *TP53Mut*-like AMLs.

*TP53Mut* AML is uniquely treatment refractory and has the worst OS among any AML subtype^1–5^. We describe molecular pathways that are commonly deregulated in this AML subtype and could offer targets for therapy. The overlap between deregulated pathways in *TP53Mut* and *TP53Mut*-like cases might suggest that both of these subsets might benefit from similar therapies. For example, we found that NFκB and EZH2 pathways are upregulated in these subsets. Both of these pathways have been implicated in AML self-renewal^37–42^. Tazemetostat, an EZH2 inhibitor, is FDA-approved and in clinical use for lymphoma^43^. Proteasome inhibitors, which are commonly used in lymphoid malignancies, attenuate the NFκB pathway^44–48^. Approaches for directly inhibiting NFκB are in clinical development^49^. A large-scale proteomic analysis of identified overexpression of MDM2, a negative regulator of p53, as a major mediator of poor prognosis in *TP53WT* AML^36^. In our study, we saw a trend of higher *MDM2* transcript levels in *TP53Mut*-like cases. Together, these findings suggest that dysfunction of the p53 pathway is a common mechanism of poor prognosis in AML. Future work could test whether targeting these pathways could offer clinical benefit in *TP53Mut* and *TP53Mut*-like AML.

Mutations in the *TP53* are largely in the DNA binding domain, but other mutation types are common as well^50^. In our study, we grouped all *TP53Mut* cases as a single entity consistent with all risk stratification schemes which define *TP53Mut* AML as adverse risk^2,25,51,52^. In AML, mutations in the *TP53* locus are most frequently missense mutations (77% missense) with frame shift (8%), nonsense (8%) and splice site (5%) accounting for the remainder of mutations^34^. Most of the missense mutations lead to loss of DNA binding and interruption of normal transcriptional regulation by p53^50^. However, missense mutations also endow neomorphic, gain-of-function transcriptional regulation by mutant p53 proteins through the interaction with novel binding partners that regulate and alter transcription^41,50,53–55^. These findings would predict that AML bearing missense *TP53Mut* might express a unique GEP relative to AML with other types of *TM53Mut*. Since the non-missense mutations account for only 33% of all *TP53Mut* cases^34^, defining the GEP of these cases would require larger sample sizes. Likewise, several groups have demonstrated that biallelic disruption of the *TP53* locus has a worse prognostic impact than single allele alterations in myeloid malignancies^30–32,56^. Future work could investigate the transcriptional differences between these alteration types in primary human AML samples.

Mutational and cytogenetic profiling are the most common molecular approaches to classify malignancies. However, the functional insights provided by transcriptional profiling has been used to reveal clinically distinct disease entities that were not detected using traditional methods. GEP of B-cell precursor acute lymphoblastic leukemia (ALL) identified as a subset of ALL that resembled Philadelphia chromosome (Ph) positive (Ph+) ALL, called Ph-like ALL. Ph-like ALLs share poor prognostic features with Ph+ ALL, including high relapse rates^19,20^. Today, Ph-like ALL is recognized as a distinct clinical entity that requires more aggressive consolidation therapy^57^. Clinical trials are ongoing to define the optimal treatment regimens for Ph-like ALL. In diffuse large B cell lymphoma, GEP revealed profiles that subdivide the disease into novel classifications, Germinal Center B (GCB) and Activated B cell large B cell lymphomas (ABC, or non-GCB)^58^. Today, GCB and non-GCB lymphomas are recognized as having unique treatment sensitivities and are managed using different approaches^59^. Recent studies captured LSC stemness-related GEP and demonstrated that a 17 gene signature could predict AML patient outcomes and be used clinically for risk stratification^17,60^. In AML, GEP can also predict therapy responses^16^. Similar to our study, patients whose *TP53WT* breast cancer expresses a GEP resembling that of *TP53Mut* breast cancer have worse OS^21^. Together, these studies demonstrate the power of transcriptional profiling to detect clinically relevant disease subtypes that are imperceptible using other methods.

Gene expression assays have provided critical tools in managing a variety of diseases^61^. Predicting cancer aggressiveness and likelihood of recurrence is an important clinical problem. Adjuvant chemotherapy is clearly beneficial in aggressive malignancies but introduces excess toxicity in patients whose tumors can be cured by surgery. Assays that detect concise gene signatures of early breast cancer provide well-validated prediction of early relapse and determine whether a patient receives adjuvant chemotherapy^62,63^. Similar assays have been developed for colon^64^ and prostate^65,66^ cancer. These assays are instrumental in optimizing therapy. For patients with ambiguous thyroid nodules, a 20 gene assay is used to determine the likelihood of invasiveness^67^. For patients with metastatic disease of unknown histology (cancer of unknown primary), a gene assay can be used to define histology^68^, which is critical for treatment selection. In heart transplant recipients, a peripheral blood gene assay is used to detect an immune signature that predicts risk of transplant rejection^69,70^. Such gene assays were developed based on large cohort GEP studies and are in widespread clinical use. Our 25 gene signatures could similarly be used to identify patients with poor prognosis who might benefit from more aggressive therapy or personalized therapy.

We find that both *TP53Mut* and *TP53Mut*-like AMLs uniquely express genes that express cell surface markers. Future work to define the cell surface protein profile of these AMLs could include these proteins. Notably, CD99, which has been proposed as a therapeutic target in AML^71^ is downregulated in both *TP53Mut* and *TP53Mut*-like cases. Once validated, the protein products of these genes could provide targets for immunotherapy or serve as labels to quickly identify these cases clinically.

Finally, our 25 signature genes could also provide insights for therapeutic targeting. This signature is expressed in both *TP53Mut* and *TP53Mut*-like patients suggesting that these targets may be effective in both patient subsets. Notably, *PEAR1* was recently found to be highly prognostic of survival in an integrated analysis of *ex vivo* drug responses and clinical outcomes of 805 patients^24^. Increased expression of *PEAR1* has been recognized in *TP53Mut* and other poor risk AML subtypes^72^. The role of *PEAR1* in leukemogenesis is not well known, but several studies have implicated this gene as a possible predictor of immune engagement in AML^72,73^. Alterations in *CREBBP* have been associated with poor prognosis in AML^74^. The t(8;16) translocation, which involves the *CREBBP* locus, is a poor prognosis abnormality included in the latest ELN recommendations for AML^25^. CREBBP is an acetyltransferase that can modify both transcription factors and histones to activate transcription^75^. CREBBP has a wide variety of binding partners implicated in leukemogenesis including p53, NFκB, hypoxia-inducible factor 1α, STAT proteins, and β-catenin^75–77^. Targeting these pathways may provide clinical benefit to this subset of patients.

These results define the unique functional attributes of *TP53Mut* AML and identify *TP53Mut*-like AML, a novel and prevalent subset of patients with similar molecular, laboratory, and clinical characteristics to *TP53Mut* AML. Our analyses suggest that *TP53Mut* and *TP53Mut*-like patients might benefit from similar treatment strategies. We find that *TP53Mut*-like AML is more prevalent than *TP53Mut* AML. Our 25 gene signature could potentially identify a cohort of patients that would benefit from *TP53Mut* AML-targeted therapies, significantly expanding the number of patients eligible for those clinical trials. Moreover, future clinical trials could be designed to target the unique molecular features of *TP53Mut* and *TP53Mut*-like AML.

---

**Supp Figure S1.**
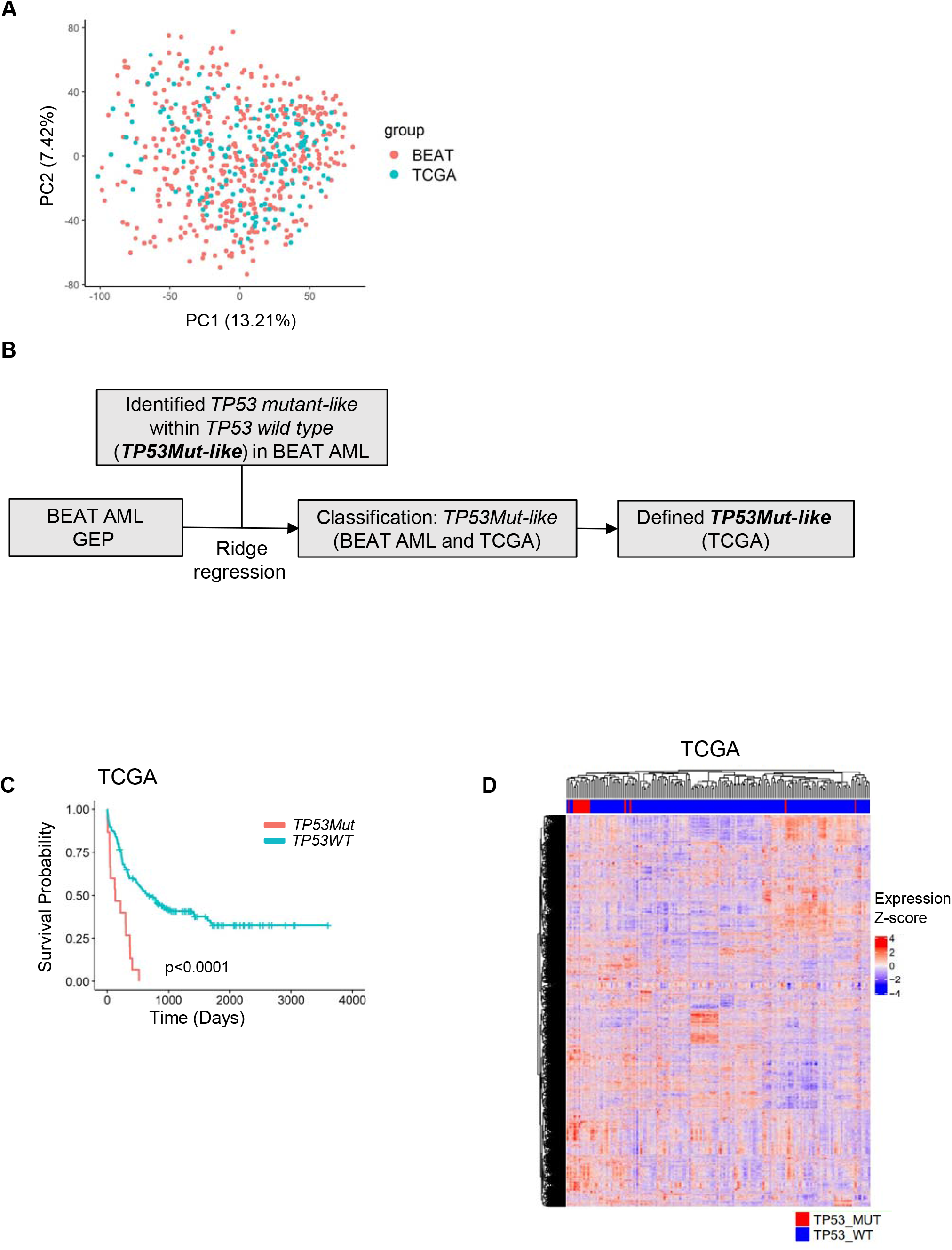

**Supp Figure S2.**
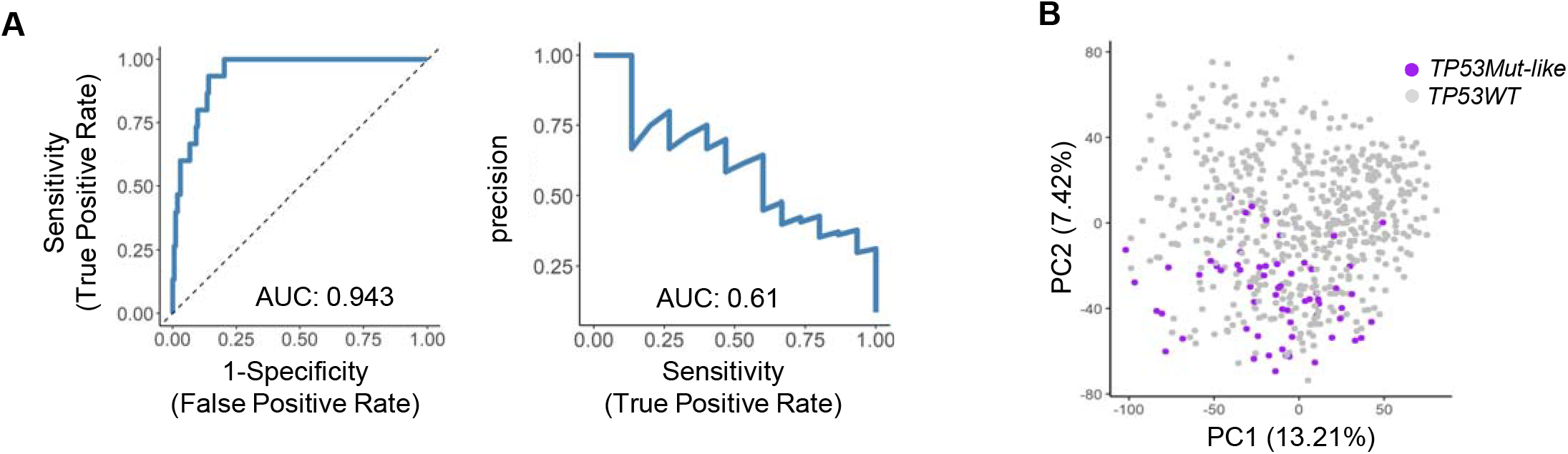

**Supp Figure S3.**
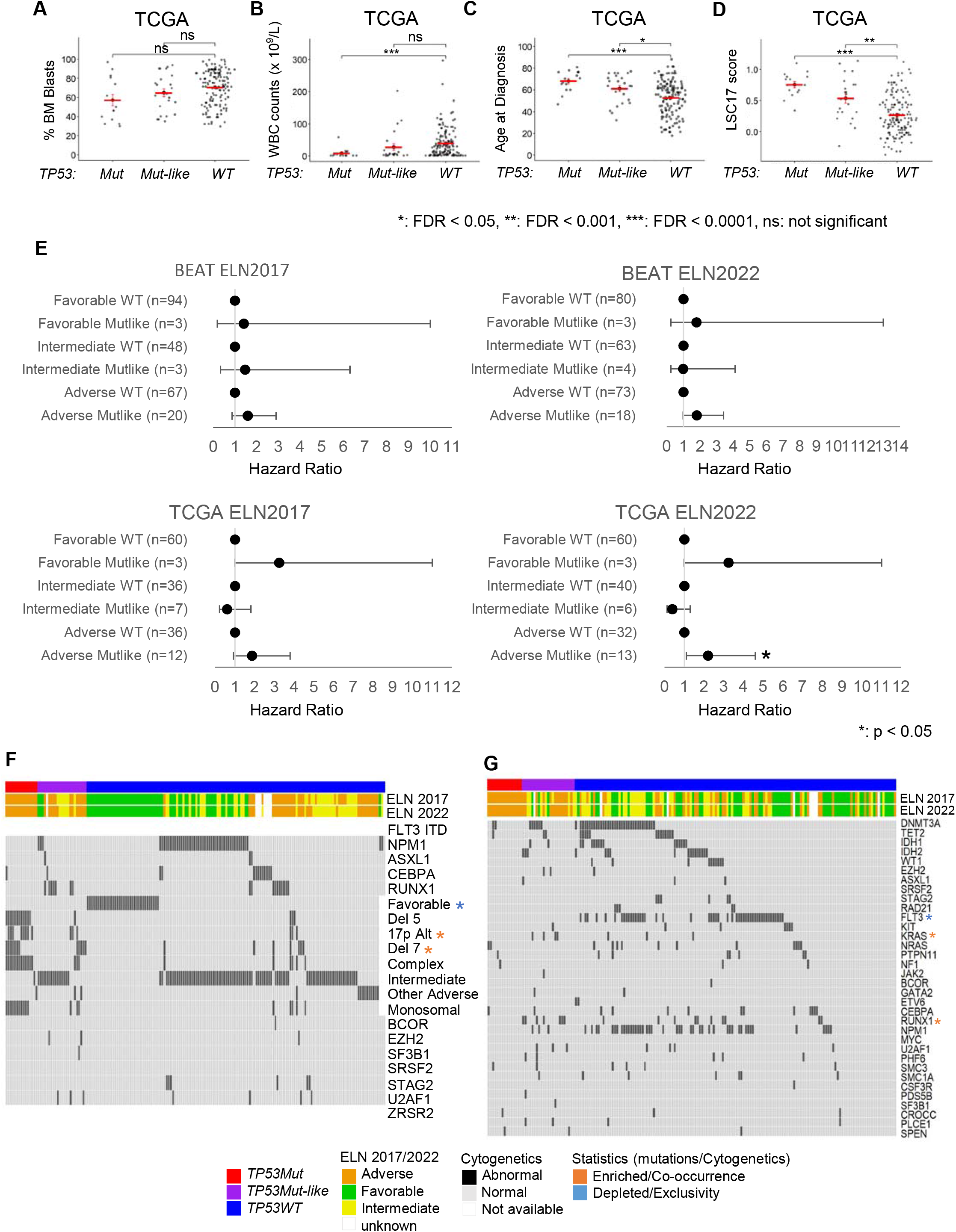

**Supplemental Figure S4.**
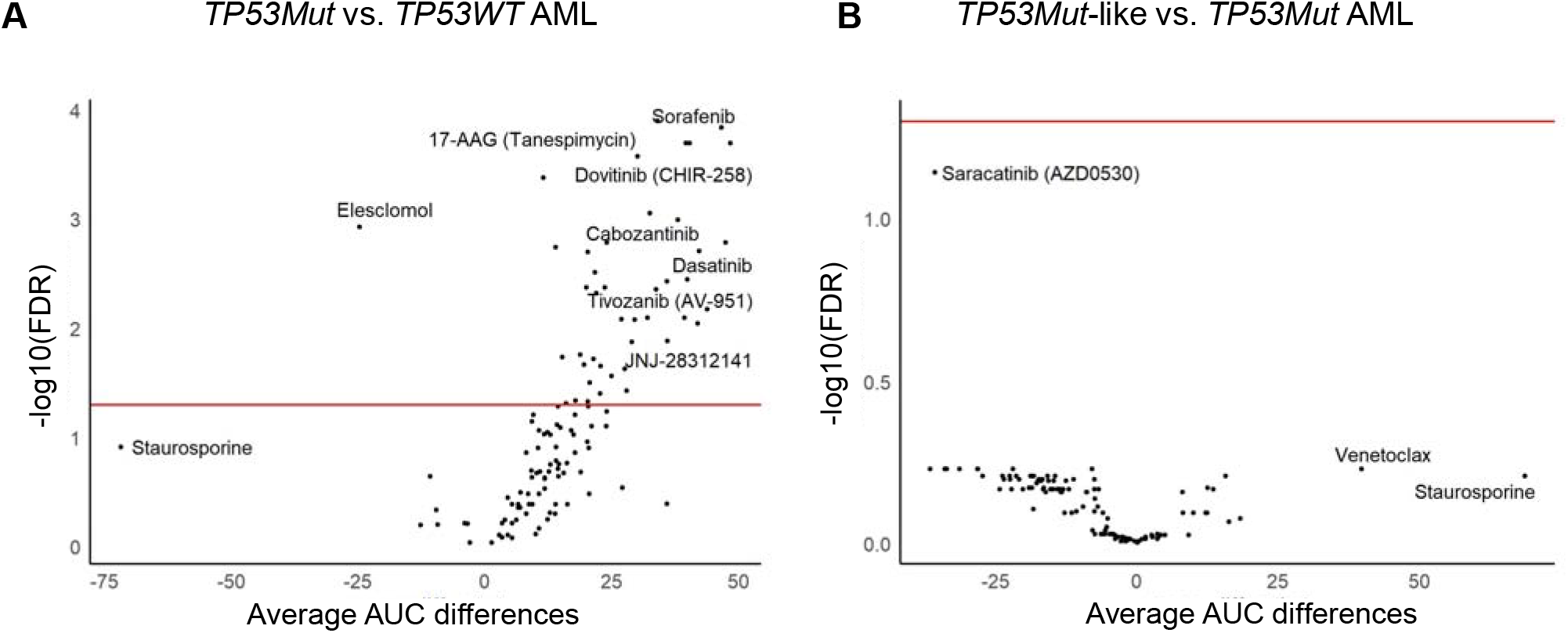
*TP53Mut*-like AML gene expression and drug sensitivity profiles

**Supplemental Figure S5.**
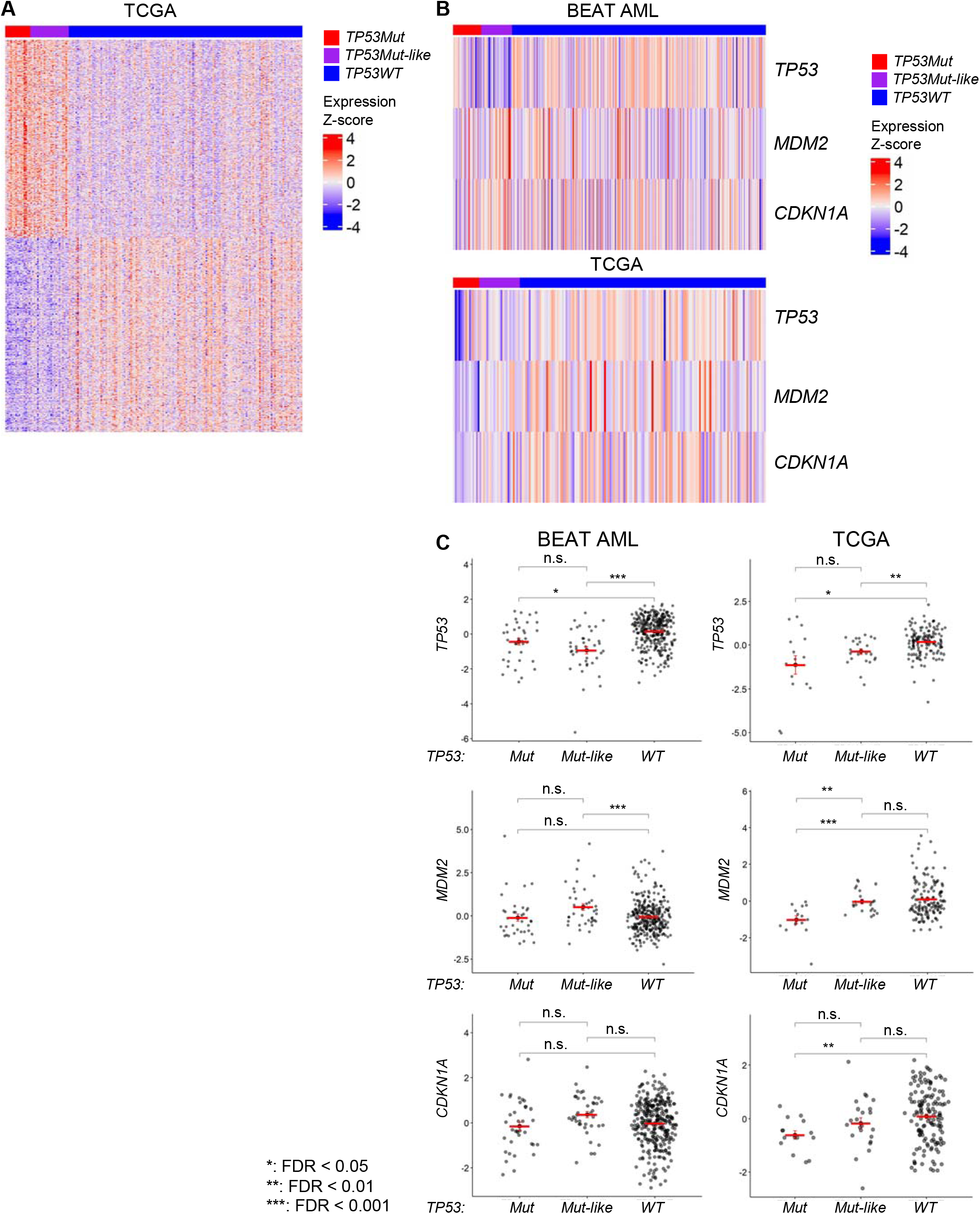
*TP53Mut*-like and *TP53Mut* AML GEP reveals unique biological pathways in TCGA

**Supplemental Figure S6.**
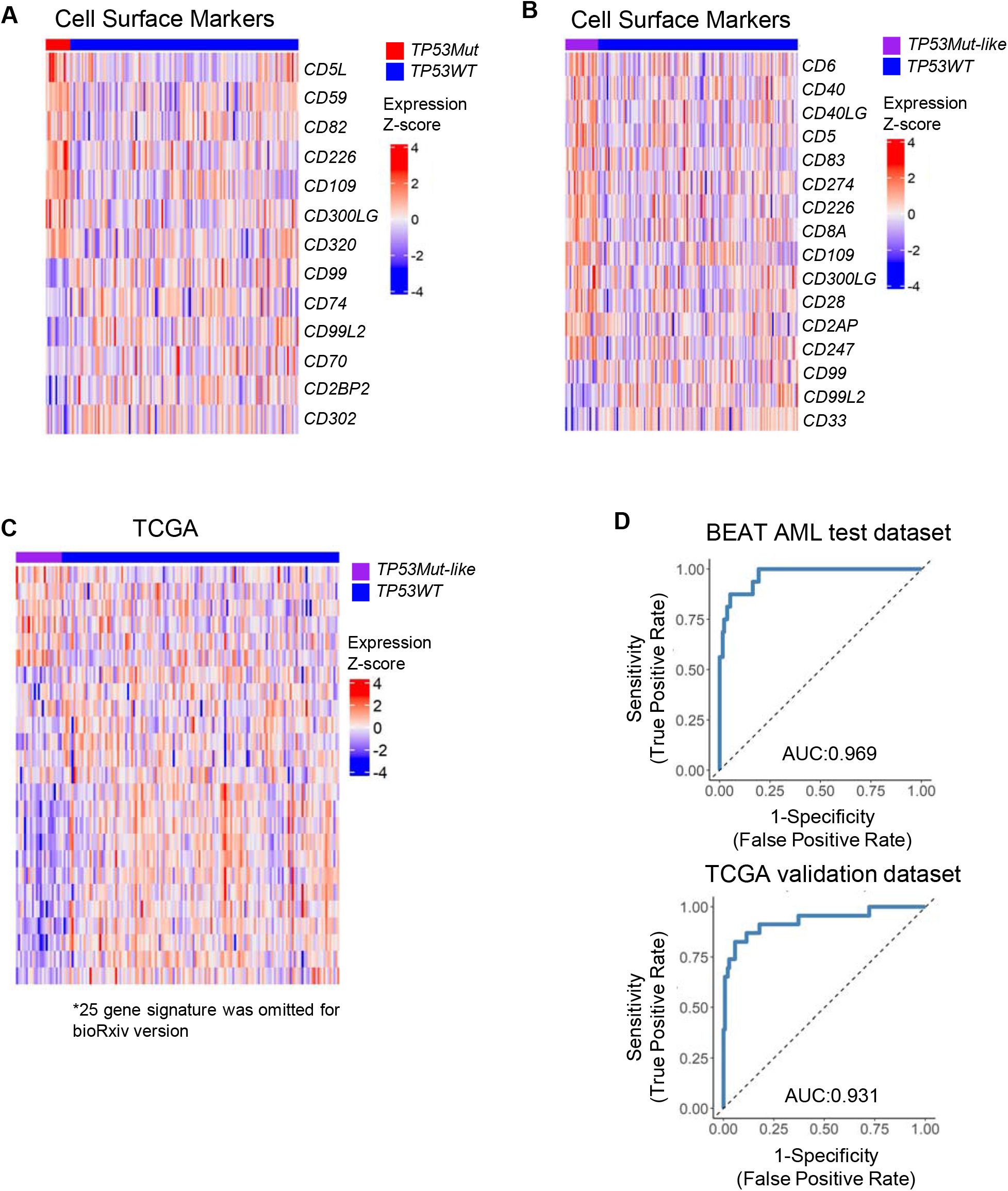
25 Signature genes define *TP53Mut*-like AML cases

